# The global biogeography of amino acid variants within a single SAR11 population is governed by natural selection

**DOI:** 10.1101/170639

**Authors:** Tom O. Delmont, Evan Kiefl, Ozsel Kilinc, Özcan C. Esen, Ismail Uysal, Michael S. Rappé, Stephen Giovannoni, A. Murat Eren

## Abstract

The diversity and geographical distribution of populations within major marine microbial lineages are largely governed by temperature and its co-variables. However, neither the mechanisms by which genomic heterogeneity emerges within a single population nor how it drives the partitioning of ecological niches are well understood. Here we took advantage of billions of metagenomic reads to study one of the most abundant and widespread microbial populations in the surface ocean. We characterized its substantial amount of genomic heterogeneity using single-amino acid variants (SAAVs), and identified systematic purifying selection and adaptive mechanisms governing non-synonymous variation within this population. Our Deep Learning analysis of SAAVs across metagenomes revealed two main ecological niches that reflect large-scale oceanic current temperatures, as well as six proteotypes demarcating finer-resolved niches. We identified significantly more protein variants in cold currents and an increased number of protein sweeps in warm currents, exposing a global pattern of alternating genomic diversity for this SAR11 population as it drifts along with surface ocean currents. Overall, the geographic partitioning of SAAVs suggests natural selection, rather than neutral evolution, is the main driver of the evolution of SAR11 in surface oceans.

## Introduction

The SAR11 family *Pelagibacteraceae* [1] represents the most ubiquitous free-living lineage of heterotrophic bacteria in the world’s oceans [2, 3, 4, 5, 6, 7]. Successful cultivation efforts have paved the way for a functional understanding of SAR11 and elucidated their critical role in marine biogeochemical cycles [8, 9, 10, 11]. Their dominance in surface seawater has resulted in great interest to understand the diversity of SAR11 and their evolution in marine habitats [12]. Environmental sequencing surveys have resolved multiple evolutionary lineages of SAR11 that possess distinct geographic distribution patterns and genomic features, and the mechanisms behind biogeographical variation across these lineages have been previously considered in the context of niche-based [13, 14] and neutral processes [15].

Despite its relevance to evolutionary dynamics and the emergence of new ecological features, to the best of our knowledge, the genome-wide diversity of individual SAR11 populations has not been characterized. A key study by Hellweger et al. [16] simulated the evolution of a few cells for large numbers of generations under a neutral model, the results of which suggested that geographic patterns in marine microbial populations could emerge without selection due to slow-moving oceanic currents. However, the hypothesis of neutral evolution as the main source of intra-population genomic variation has not been empirically tested.

Here we took advantage of successful cultivation efforts, whole genome sequences, and recent environmental metagenomic surveys to characterize the genomic diversity of a single SAR11 population across distant geographies, herein named ‘SAR11 Low-Latitude Population A’ (S-LLPA). This population alone recruited more than one percent of surface ocean metagenomic reads worldwide and might be the most widespread and abundant microbial population in low-latitude surface oceans. We developed a novel approach to characterize single-amino acid variants (SAAVs) from metagenomic data and investigated evolutionary dynamics operating on S-LLPA. Our analysis of SAAVs revealed multiple subpopulations within S-LLPA, whose distributions were linked to large-scale oceanic current temperatures. Our empirical data revealed a stronger emphasis on natural selection, rather than neutral evolution, as the driver of the evolution of SAR11 in low-latitude surface oceans.

## Results

We characterized the relative abundances of 21 SAR11 isolates across four oceans and two seas using a dataset of 103 metagenomes, including 93 samples from the TARA Oceans Project [17] and 10 samples from the Ocean Sampling Day Project [18] to cover high-latitude areas of the Northern hemisphere. All metagenomes corresponded to small planktonic cells (0.2-3m in size) from the surface (0-15 meters depth; n=71) and deep chlorophyll maximum (17-95 meters depth; n=32) layers of the water column (Table S1). The SAR11 isolates we used belonged to subclades Ia.1 (n=6), Ia.3 (n=11), II (n=1), IIIa (n=2) and Va (n=1) (Table S1), which collectively recruited 1,029,716,339 reads from all metagenomes and corresponded to 3.3% of the dataset (Table S2).

### The metapangenome of SAR11

To investigate large-scale links between genomic traits and niche partitioning of SAR11 populations, we performed a pangenomic analysis in conjunction with read recruitment from the metagenomic data. The pangenome of the 21 SAR11 genomes consisted of all 29,719 genes grouped into 6,175 protein clusters (PCs) (Table S3). The clustering of genomes based on shared PCs matched that of the previously described phylogenetic clades [19] (Figure 1). The distribution of the SAR11 pangenome across metagenomes, which we define as the metapangenome, revealed distinct distribution patterns for each clade within SAR11 (Figure 1). Clade Ia recruited the most reads compared to other clades (Table S2), consistent with previous studies that found this clade to be highly abundant in surface waters [15, 13, 14]. Pangenomic traits divided clade Ia into two main clusters, which correspond to the high-latitude subclade Ia.1 and the low-latitude one, Ia.3 (Figure 1). While all high-latitude genomes displayed a bi-polar geographic distribution in the metagenomic dataset, low-latitude genomes were clustered into multiple sub-groups with different patterns of geographic distribution, suggesting further refinement of subclade designations may be possible. Nevertheless, our analysis exposed a strong agreement between the phylogeny, pangenome, and ecological niche partitioning of members in the SAR11 clade Ia (Figure 1).

**Figure 1.**
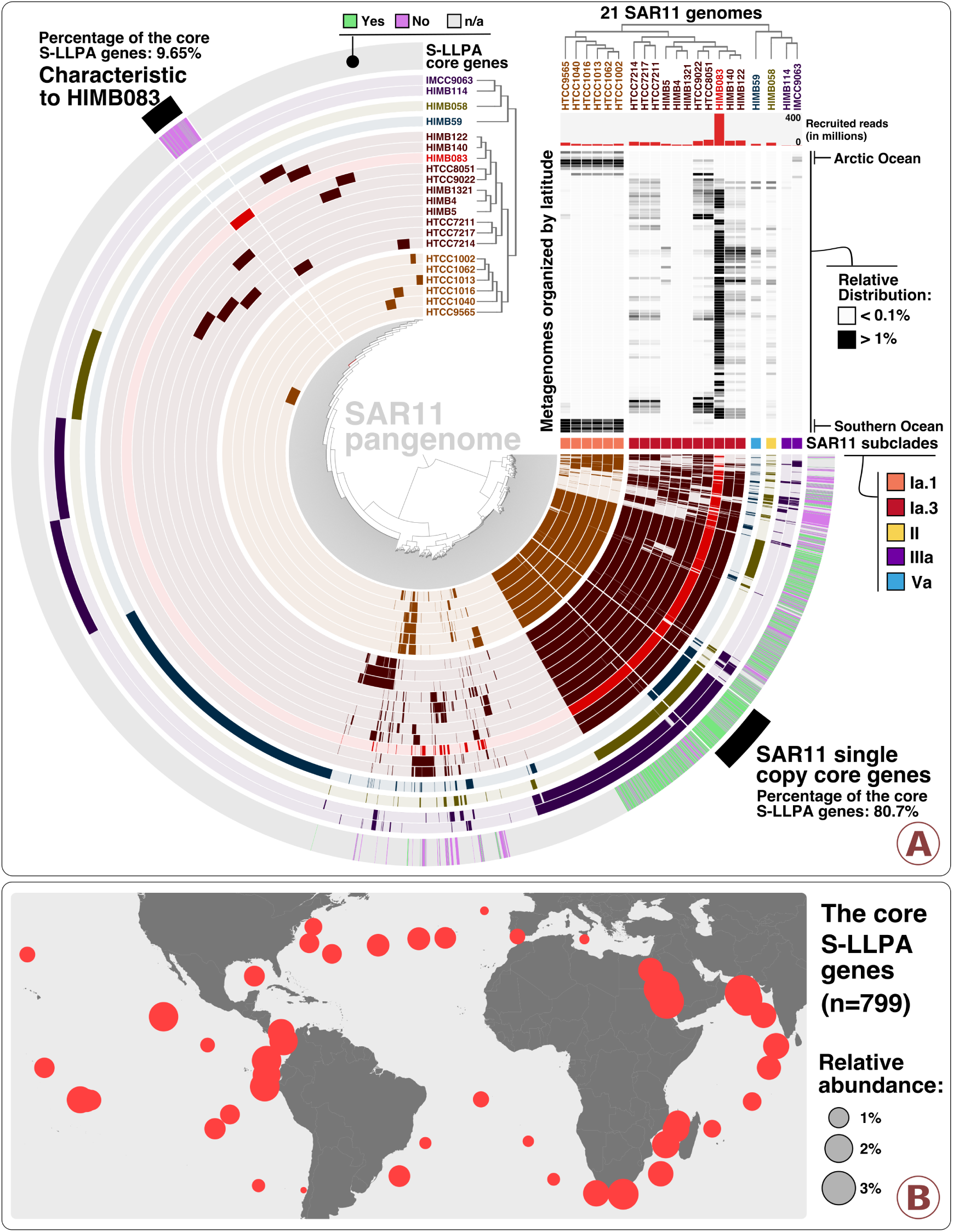
Linkage between the pangenomic traits and large-scale distribution of SAR11 genomes. Panel A, interactive version of which is also available at the URL https://anvi-server.org/merenlab/sar11_metapangenome, describes the metapangenome of 21 SAR11 isolate genomes based on the occurrence of 6,175 protein clusters, in conjunction with their phylogeny (clade level) and relative distributions in 103 metagenomes ordered by latitude from the North Pole to the South Pole (top right heat map). The relative distributions were displayed for a minimum value of 0.1% and a maximum value of 1%. Protein clusters associated with HIMB83 were colored based on the occurrence of the 799 core genes of S-LLPA. Panel B describes the relative abundance of the 799 core genes of S-LLPA across surface metagenomes from TARA Oceans.

### A remarkably abundant and widespread SAR11 population at low latitudes

Although the low-latitude subclade Ia.3 was overall the most abundant SAR11 lineage in the dataset, genomes within this group differed remarkably in their competitive recruitment of short reads (Figure 1, Table S2). For example, while the least abundant Ia.3 genome, HIMB1321, recruited only 13.2 million reads, the most abundant one, HIMB83, recruited 390.9 million reads. This is 47.3% of all reads recruited by the eleven Ia.3 genomes and 1.18% of the entire metagenomic dataset (Table S2). HIMB83, isolated from coastal seawaters off Hawai’i, USA, has a 1.4 Mbp genome, 1,470 genes, and a GC-content of 29.1%. HIMB83 recruited two times more reads than the most abundant *Prochlorococcus* isolate genome from the same set of metagenomic data (Table S2), and nearly 10X more reads than the most abundant metagenome-assembled genome reconstructed de novo from the TARA Oceans metagenomes [20] (Table S2). Its remarkable abundance has also been recognized by others [21, 22]. To improve readability, we named this widespread and abundant SAR11 population we accessed through HIMB83 as the ‘SAR11 Low Latitude Population A’ (S-LLPA). To the best of our knowledge, S-LLPA is the most abundant and widespread microbial population in the euphotic zone of low-latitude oceans and seas.

As a single member of the S-LLPA, the genomic context HIMB83 provides is not exhaustive. However, it gives access to the core S-LLPA genes through read recruitment. To identify core S-LLPA genes we used a conservative two-step filtering approach. First, we defined a subset of the 103 metagenomes as the main ecological niche of S-LLPA using genomic mean coverage values (Table S2, supplemental information). Our selection of 74 metagenomes in which the mean coverage of HIMB83 was >50X encompassed three oceans and two seas between -35.2° and +43.7° latitude, and temperatures between 14.1°C and 30.5°C (Figure S1, Table S1). From these 74 metagenomes, we then defined a subset of HIMB83 genes as the core S-LLPA genes if in each of the 74 metagenomes, the base-5 logarithm of the mean coverage of a gene remained within ±1 of that for the mean coverage across all genes. Figure S1 displays the coverage of all HIMB83 genes across all metagenomes, and Table S4 reports the coverage statistics. While the 799 genes that met this criterion systematically occurred within the niche boundaries of S-LLPA, most of the 671 genes filtered out coincided with hyper-variable genomic loci (Table S4) and occurred mostly in PCs unique to HIMB83 within the SAR11 metapangenome (Figure 1). Thus, the metapangenome revealed that genes unique to HIMB83 were not commonly found in S-LLPA. Hyper-variable genome regions are common features of surface ocean microbes [23, 24, 25] that are not readily addressed through metagenomic read recruitment. On the other hand, the 799 core S-LLPA genes (defined by coverage) were strongly associated with PCs comprising the SAR11 Ia core-genome (defined by sequence similarity), suggesting they cover a large fraction of the S-LLPA genomic backbone (Figure 1). Core S-LLPA genes recruited on average 1.25% of reads in the 74 metagenomes (Figure 1, Table S4). Not only does their broad geographic prevalence suggest dispersion is not a limiting factor for SAR11 in surface seawater of low-latitude oceans and seas, but that these properties of S-LLPA also present a unique opportunity to study the genome-wide diversity and evolutionary dynamics of a single marine microbial population across distant geographies.

### Single-nucleotide variants were widespread across the core S-LLPA genes

To investigate the genomic diversity of S-LLPA within the boundaries of its niche, we first characterized all single-nucleotide variants (SNVs) in the 799 core genes. Our analysis revealed a total of 10,807,040 SNVs across the 74 metagenomes (Table S2). 80.5% of all SNVs occurred in the third nucleotide position of codons, while the remaining SNVs were shared unevenly between the first (14%) and second (5.5%) nucleotide positions (Table S2). The density of SNVs (i.e. the percentage of SNVs per hundred nucleotides of DNA) were similar across the 74 metagenomes (Figure S1) and averaged to 19.3% across all core S-LLPA genes, revealing substantial genomic diversity within this population. SNV density for each gene individually ranged from as low as 2.9% (a gene coding a cold-shock DNA-binding protein in the Indian Ocean) to as high as 37.3% (a gene of unknown function in the Pacific Ocean) (Figure S1 and Table S5), indicating the variedness of diversification across core S-LLPA genes.

### SNVs to SAAVs: Metagenomic read frequencies resolve variation for codons containing multiple SNVs

To study genomic variation that impacts amino acid sequence, we implemented a framework to characterize amino acid substitutions in metagenomic data (see Materials and Methods). Briefly, our approach employs metagenomic short reads covering all three nucleotides in a given codon to determine the frequency of single-amino acid variants (SAAVs) in translated protein sequences. The use of short reads mapping entirely to a given codon position preserves the linkage between variable nucleotides. Quantifying frequencies of identical reads in a given codon position allowed us to report accurate estimations of non-synonymous variation for codons containing SNVs in more than one position, addressing a fundamental problem first introduced three decades ago [26, 27]. This was essential for our survey of non-synonymous variation in S-LLPA core genes, as on average 22.5% of SNVs per metagenome co-occurred with other SNVs in the same codon.

Among the 799 core S-LLPA genes and 74 metagenomes, we identified 1,074,096 SAAVs in which >10% of amino acids diverged from the consensus (i.e. the most frequent amino acid for a given metagenome) (Table S2). The average SAAV density across core genes was 5.76% and correlated with SNV density (R^2^=0.89) (Figure S1). To improve downstream beta-diversity analyses, we discarded codon positions if their coverage in any of the 74 metagenomes were <20X, which resulted in a final collection of 738,324 SAAVs across 37,416 amino acid positions (14.8% of all codons) within the core genes (Table S6). Table S5 reports SNV and SAAV densities for each core S-LLPA gene across all metagenomes after applying the 20X minimum coverage cut-off. All genes except a 679 nt long ABC transporter contained at least one SAAV in at least one metagenome, revealing a wide range of amino acid sequence diversification traits among core S-LLPA proteins. Evidence for protein sweeps (i.e. the removal of sequence diversity by strong positive selection for a single protein variant) was rare. We consider a protein has undergone a sweep in a given environment if >90% of the reads recruited from the metagenome resolve to a single amino acid residue for every codon position in the corresponding gene (i.e. the gene lacks SAAVs).

### Purifying selection governs the identity of amino acid substitution types

Each SAAV is a vector of 21 items: frequencies of the 20 amino acids and the stop codon in a metagenome. SAAVs were often dominated by a few amino acids, resulting in a large degree of sparsity. To study patterns of amino acid variability, we simplified the SAAV data by associating each SAAV position with an amino acid substitution type (AAST), defined as the two most frequent amino acids found in a given SAAV. In the 738,324 SAAVs, we observed 182 of 210 theoretically possible unique AASTs. The distribution of AASTs was highly consistent across metagenomes, regardless of geographic location (Figure 2, Table S7). We used the BLOcks Substitution Matrix 90 [29] (BLOSUM90) as a proxy to assess the functional difference between the pairs of amino acids in each AAST with scores ranging from ‘-6’ (highly dissimilar) to ‘+3’ (highly similar). ‘Isoleucine/valine’ was the most commonly found AAST in our study and represented more than 10% of all SAAVs (Figure 2). This is not surprising given that ‘isoleucine/valine’ has a maximum BLOSUM90 score, indicating that valine ↔ isoleucine substitutions are statistically less likely to disrupt protein stability and therefore more likely to represent permissible, functionally conserved substitutions. In general, prevalent AASTs were biased towards higher BLOSUM90 scores (Table S7). For instance, the BLOSUM90 scores for the 10 most and least prevalent AASTs averaged to +0.9 and -3.71, respectively, indicating that the majority of SAAVs likely represent functionally conserved substitutions. It furthermore suggests a powerful influence of purifying selection via the deletion of mutations disruptive to protein architecture. That said, the BLOSUM90-defined chemical similarity between amino acids was not the sole predictor of the abundance of AASTs in our dataset (R^2^: 0.3) (Figure 2, Table S7). For instance, the AAST ‘isoleucine/threonine’ (isoleucine is hydrophobic and threonine has an alcohol functional group that can participate in hydrogen bonding) was the most prevalent substitution with a negative BLOSUM90. This unlikely substitution represented 2.92% of all SAAVs. Another widely used substitution matrix, BLOSUM62 [29], mirrored these trends (Table S7). Overall, 20.8% of SAAVs exhibited non-conservative substitutions as indicated by negative BLOSUM90 scores, suggesting a role for adaptive processes for functional diversification of S-LLPA proteins. These AASTs are candidates for diversifying selection, which in principle could yield multiple phenotypes within the global S-LLPA population.

**Figure 2.**
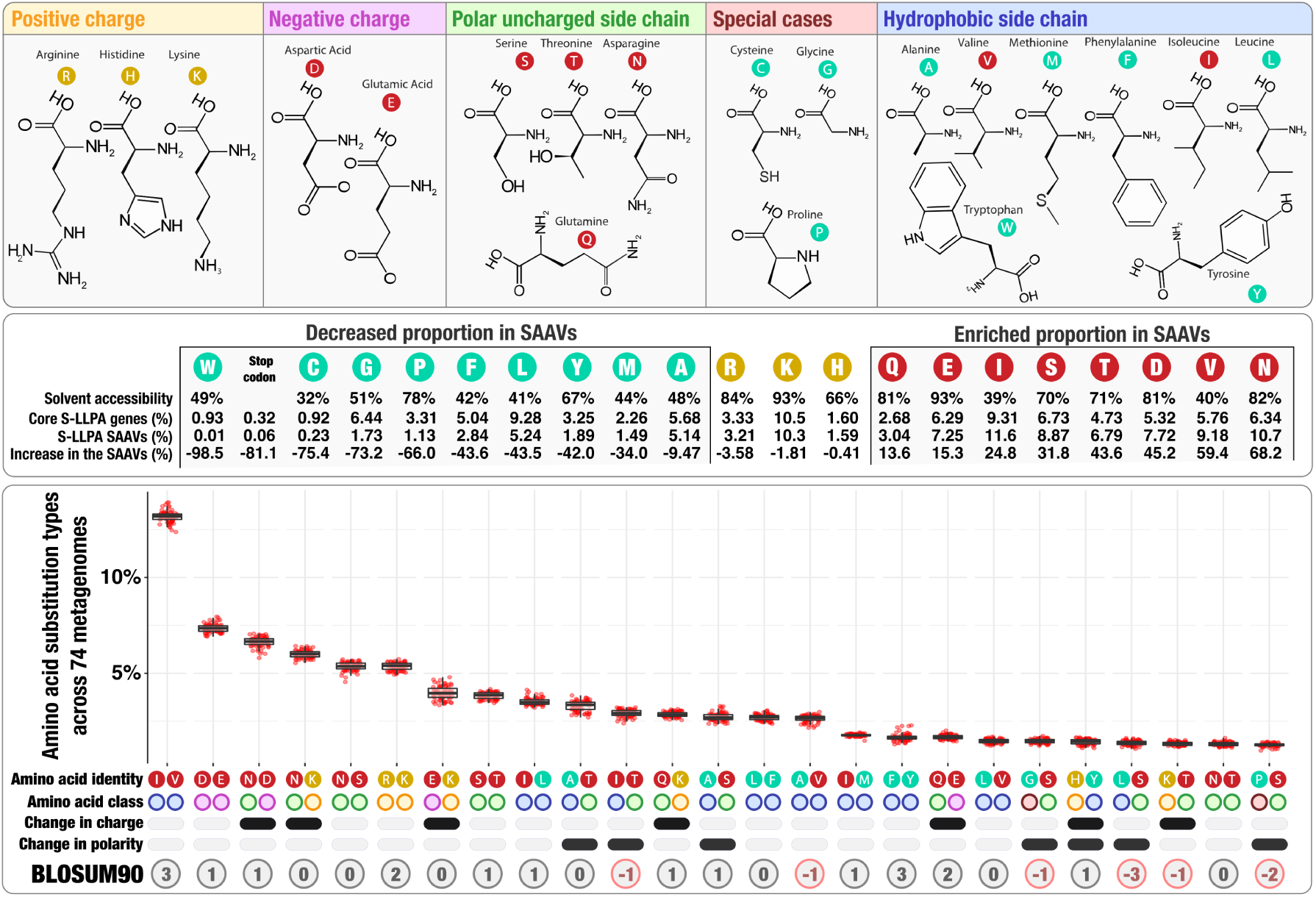
Physico-chemical properties of amino acid variants. The top panel describes the structure of 20 amino acids grouped by their main chemical properties. The middle panel describes the solvent accessibility of amino acids, and their relative distribution in the core S-LLPA genes and SAAVs. The bottom panel describes (1) the relative abundance of amino acid substitution types (AASTs) across 74 metagenomes (boxplots), (2) classes of amino acids each AAST contains, and (3) the BLOSUM90 score for amino acids in each AAST. AASTs included in the analysis occur in more than 1% of SAAVs (n=25; corresponds to 87.1% of all SAAVs). The solvent accessibility of amino acids derives from the analysis of 55 proteins [28].

The occurrence of amino acids in SAAVs did not strongly correlate with their occurrence in core S-LLPA genes (R^2^: 0.65) (Figure 2). All amino acids with negative charges (Asp, Glu) and polar amino acids with uncharged side chains (Thr, Asn, Ser, Gln) were enriched in SAAVs compared to the overall nucleotide composition of the core S-LLPA genes (Figure 2). For instance, while asparagine made up 6.34% of all amino acids in the core genes, 10.7% (±0.16%) of SAAVs involved asparagine substitutions across the 74 metagenomes (Table S7). In contrast, amino acids with positive charges did not exhibit substantial differences (<4% variation). Finally, our data showed that, except for valine and isoleucine, all amino acids with a hydrophobic side chain were less represented in SAAVs (Figure 2, Table S7). Amino acids that form the hydrophobic core of proteins (with low solvent accessibility) are critical for their stability and mutations in hydrophobic sites are often deleterious [30, 31]. Consequently, solvent accessibility positively correlates with amino acid polymorphisms in bacterial models [32]. Within S-LLPA, solvent accessibility largely explained the differential involvement of amino acids in SAAVs, revealing (1) a strong correspondence between functional classes of amino acids and their stability in the environment, and (2) a systematic influence of purifying selection against deleterious variants in the hydrophobic core of S-LLPA proteins.

### The geographic partitioning of SAAVs reflects oceanic current temperatures

We used Deep Learning (see Material and Methods) to estimate relationships between metagenomes based on our curated collection of 738,324 SAAVs (Table S8). Hierarchical clustering of samples based on Deep Learning-estimated distances resulted in two main groups (Figure 3). The first group (n=33) encompassed all metagenomes from the Red Sea and Indian Ocean, as well as metagenomes from the West side of the Atlantic Ocean (Figure 3). The second group (n=41) encompassed most metagenomes from the Pacific Ocean, as well as the East side of the Atlantic Ocean (except near the southern tip of Africa) and the Mediterranean Sea. Thus, the Atlantic Ocean was partitioned according to longitude rather than latitude. Samples collected from the deep chlorophyll maximum layer of the water column mirrored trends observed in the surface samples (Figure S2). Interestingly, the distribution of samples in these groups reflected ocean current temperatures at large scale: while the first group was associated with warm currents (Agulhas, Mozambique, Brazil and Gulf stream), the second group was associated with cold currents (Benguela, Canary, California and Peru). The association between SAAVs and ocean current temperatures revealed a strong, global signal at the amino acid-level for S-LLPA’s response to changing temperatures and suggested the presence of two main ecological niches for this prevalent population. Interestingly, the niche defined by cold currents exhibited significantly more SAAVs (ANOVA, p:1.66e-12) and a significantly higher SAAVs/SNVs ratio (ANOVA, p:1.07e-10). This observation could be explained either by (1) extinction/re-emergence events that operate on similar amino acid positions, or (2) changes in abundances within a large seed bank of variants due to positive selection as the population moves through currents. Although testing these hypotheses requires more suitable experiments, a neutral evolution hypothesis in unable to account for this phenomenon.

**Figure 3.**
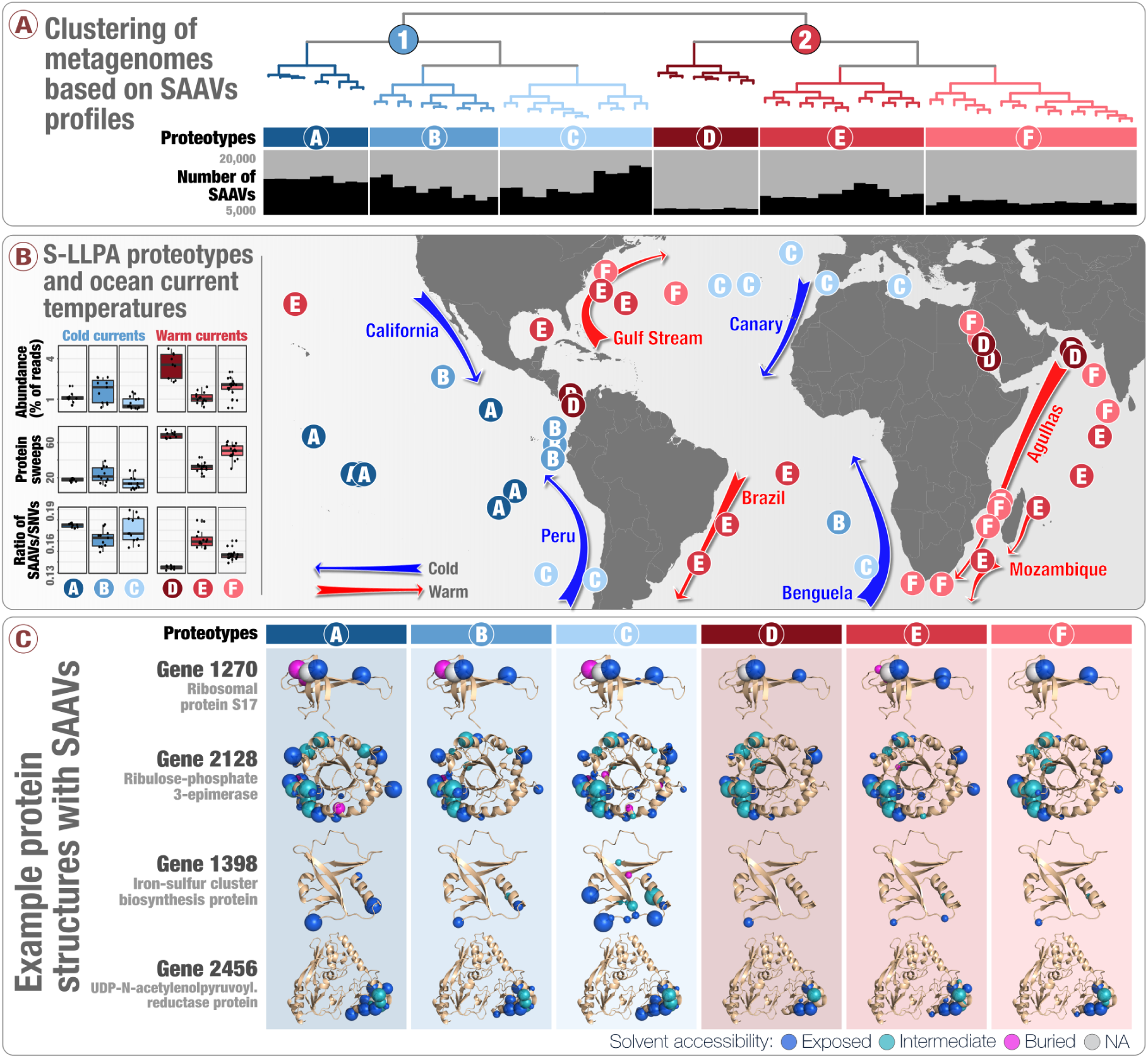
Metagenomic clustering and geographic partitioning of S-LLPA main groups and proteotypes at large scale. Panel A describes the organization of 74 metagenomes based on SAAV data. The world map in panel B displays the geographic partitioning of the two main metagenomic groups and six proteotypes. Panel B also describes the relative abundance of S-LLPA, number of protein sweeps and the SAAVs to SNVs ratio across the six proteotypes. Panel C displays SAAV positions on the predicted protein structures of four core S-LLPA genes across six proteotypes. Size of a SAAV is positively correlated with their prevalence (from 0% to 100%) in metagenomes in the same proteotype.

To explore more detailed trends unexplainable by ocean current temperatures alone, we further divided our dendrogram into six sub-clusters based on the elbow of the intra-cluster sum-of-squares curve of k-means clusters (Figure S3). These ‘S-LLPA proteotypes’ grouped samples with similar amino acid diversification traits (Figure 3). While the temperature at the time of sampling was not an insignificant predictor of the groupings (ANOVA, p:3.59e-07), the number of SAAVs, the number of protein sweeps, and the SAAVs/SNVs ratio were more significant predictors (ANOVA, p:<2e-16). Most S-LLPA proteotypes linked samples from distant geographical regions (Figure 3). An exception to this was the proteotype A, which only contained Pacific Ocean metagenomes (Figure 3). For instance, proteotypes E and F occurred both in the Indian Ocean and the West side of the Atlantic Ocean and associated with distinct warm currents: E was characteristic of the Mozambique and Brazil currents while F dominated the Agulhas current (Figure 3). One of the most interesting proteotypes, D, contained a distinctively low number of SAAVs and SAAVs/SNVs ratio, despite its high relative population abundance, and grouped metagenomes sampled from both sides of the Panama Canal with metagenomes from the Red Sea and North of the Indian Ocean (Figure 3). Our analysis suggests that similar environmental pressures in these locations may have resulted in increased protein sweep events in similar genes in S-LLPA (Table S5). Overall, both the main ecological niches and more refined proteotypes suggest that amino acid diversification traits could link distant geographic regions, conflicting with a simulation based on a neutral evolution model [16] (Figure S4).

### A nexus between protein structure properties and the occurrence of SAAVs

From overall trends, we finally turned our attention to SAAVs in the context of protein structures. To visualize the relationship between the two, we mapped SAAVs from the core S-LLPA genes onto predicted protein structures for 436 of the core S-LLPA genes yielding a match in the Protein Data Bank (PDB) [33]. We found that buried amino acids (i.e. those with low solvent accessibility) contained far fewer SAAVs (ANOVA, p:<2e-16), an observation strikingly apparent in TIM barrels (Figure 3; also see http://anvio.org/data/S-LLPA-SAAVs/), where SAAVs mostly occur in the solvent accessible outer regions. Homologous protein families have decreased variability in their hydrophobic cores, and our study corroborates these findings within a naturally occurring population. Moreover, solvent inaccessible SAAVs exhibited lower BLOSUM90 scores compared to solvent accessible ones (ANOVA, p:<2e-16) (Table S9), suggesting they likely alter protein architecture. In addition, we identified domain-specific SAAV signatures across geographies (Figure S5), indicating that domains in a single SAR11 protein are not necessarily synchronized in their response to environmental changes. Overall, these observations suggest that the nexus between SAAVs and predicted protein structures can reveal variants of known proteins persistent in the environment, and offer an avenue for structural biologists to gain additional insights into protein evolution through metagenomics.

## Discussion

We took advantage of billions of metagenomic reads to investigate single-amino acid variants (SAAVs) in a single SAR11 population remarkably abundant across temperate and tropical latitudes. The results posit a powerful argument for purifying selection constraining the scope of neutral evolution to amino acid sequence variants permitted by protein stability requirements. Permissible variation revealed geographically bounded amino acid diversification traits, the partitioning of which reflect large-scale ocean current temperatures. Previous studies have subdivided SAR11 clade Ia into cold-water (Ia.1) and warm-water (Ia.3) subclades with distinct latitudinal distributions [13], and reported sinusoidal oscillations between their abundances as a function of seawater temperature at a single temperate ocean site [14]. Our findings suggest a similar influence of temperature on SAR11 at much finer scales of evolution, supporting the perspective that temperature and its co-variables drive plankton diversification across multiple scales of evolution. Significantly more protein variants in cold currents and more frequent protein sweeps in warm currents reveal a global pattern of alternating diversity for the S-LLPA as it drifts along with surface ocean currents in temperate and tropical latitudes.

Our findings resonate well with recent observations on another abundant and prevalent marine lineage, *Prochlorococcus.* In a key study using high-throughput single-cell genomes isolated from the same site over time, Kashtan et al. [23] identified subpopulations containing distinct ‘genomic backbones’, with each backbone exposing a unique set of accessory genes. If ‘genomic backbones’ within SAR11 populations echo those observed in *Prochlorococcus*, then accessory genes associated with the subpopulations we have identified are likely central to the population’s fitness. Moving forward, combining high-throughput cultivation and single-cell genomics with metagenomics from distant geographies could fill a critical gap between core gene variations and the ecology of accessory genes in the context of oceanic currents and other key environmental variables.

## Materials and Methods

The URL http://merenlab.org/data/2017_Delmont_et_al_SAR11_SAAVs/ contains a reproducible work flow that extends the descriptions and parameters of programs used here for (1) the metapangenome of SAR11 using cultivar genomes, (2) the profiling of metagenomic reads the cultivar genomes recruited, (3) the analysis of single nucleotide variants using Deep Learning, and (4) the visualization of single nucleotide variants in the context of protein structures. For all anvi’o analyses, we used the anvi’o v2.4.0 (available from http://merenlab.org/software/anvio/).

### SAR11 cultivar genomes

We acquired the genomic content of 21 SAR11 isolates from NCBI and simplified the deflines using anvi’o [34]. We then concatenated all contigs into a single FASTA file, and generated an anvi’o contigs database, during which Prodigal [35] v2.6.3 identified open reading frames in contigs, and we annotated them with InterProScan [36] v1.17. Table S1 reports the main genomic features.

### Metagenomic datasets

We acquired 103 metagenomes from the European Bioinformatics Institute (EBI) repository under the project ID ERP001736 (n=93; TARA Oceans project) and ID ERP009703 (n=10; Ocean Sampling Day project), and removed noisy reads with the illumina-utils library [37] v1.4.1 (available from https://github.com/meren/illumina-utils) using the program ‘iu-filter-quality-minoche’ with default parameters, which implements the method previously described by Minoche et al [38]. Table S1 reports accession numbers and additional information (including the number of reads and environmental metadata) for each metagenome.

### Pangenomic analysis

We used the anvi’o pangenomic workflow to organize translated protein sequences from SAR11 genomes into protein clusters. Briefly, anvi’o uses BLAST [39] to assess the similarity between each pair of amino acid sequences among all genomes, and then resolves this graph into protein clusters using the Markov Cluster algorithm [40]. We built the protein clustering metric using a minimum percent identity of 30%, an inflammation value of 2, and a maxbit score of 0.5 for high sensitivity. Table S3 reports the occurrence of protein clusters across genomes. We clustered both the SAR11 genomes and protein clusters based on the pangenomic metric using Euclidian distance and Ward ordination.

### Recruitment and profiling of metagenomic reads

We mapped reads from each metagenome against the single FASTA file containing the collection of SAR11 genomes using BOWTIE2 [41] v.2.0.5 with default parameters, and stored the results as BAM files using SAMTOOLS [42] v1.3.1. To confirm our observations, we also used BWA [43] to recruit reads (with the option n=0.05). We used anvi’o to generate profile databases from the BAM files and combine these mapping profiles into a merged profile database, which stored coverage and variability statistics as outlined in Eren et al [34]. Table S2 reports the mapping results (number of recruited reads, mean coverage, detection) across the 103 metagenomes.

### Determining the coverage of HIMB83 genes across metagenomes

The merged profile database contains the detection of individual genes across metagenomes. We normalized the detection of HIMB83 genes in each metagenome (summarized in the Table S4) and calculated their coefficient of gene variation. We used it to identify metagenomes in which HIMB083 was well detected but with a high instability in genes coverage, indicative of non-specific reads recruitment.

### Determining the main ecological niche and core genes of S-LLPA

We considered metagenomes in which HIMB83 was sufficiently detected (mean genomic coverage >50X) with a stable genes level detection (coefficient of gene variation <1.25) to represent the main ecological niche of S-LLPA (see supplemental information for more details). The 74 metagenomes fitting these criteria are summarized in the Table S1. We then considered HIMB83 genes to be sufficiently linked to S-LLPA when their mean coverage never diverged more than five times from the average mean coverage of all genes across the 74 metagenomes. The 799 genes fitting this criterion are summarized in the Table S4.

### Generating single-nucleotide variants (SNV) data

We used the program ‘anvi-gen-variability-profile’ to report variability tables describing the nucleotide frequency (i.e., ratio of the four nucleotides) in recruited metagenomic reads per SNV position. To study the extent of variation of the core S-LLPA genes across all metagenomes, we instructed anvi’o to report positions with more than 1% variation at the nucleotide level (i.e., at least 1% of recruited reads differ from the consensus nucleotide). To determine the SAAV to SNV ratio (described below), we instructed anvi’o to report only positions with more than 10% variation at the nucleotide level. Table S2 reports the density of SNVs for all SAR11 genome across all metagenomes. We also used anvi’o to report SNVs for a subset of genes and metagenomes, and by considering only nucleotide positions with a minimum coverage cut-off across metagenomes under consideration. Controlling the minimum coverage of single nucleotide positions across metagenomes improves confidence in variability analyses. Table S5 reports the SNV density values for all core S-LLPA genes.

### Generating single-amino acid variants (SAAVs) data

The program ‘anvi-gen-variability-profile’ (with an additional ‘–engine AA’ flag) reported variability tables describing the single-amino acid frequency data (i.e., frequency counts of the 20 amino acids plus the stop codon) per codon position. Anvi’o only considers short reads that cover the entire codon to determine amino acid frequency at a given codon position in a metagenome. We instructed anvi’o to report only positions with more than 10% variation at the amino acid-level (i.e., at least 10% of recruited reads differ from the consensus codon). Table S2 reports the density of SAAVs for all SAR11 genome across all metagenomes. We also used anvi’o to report SAAVs for a subset of genes and metagenomes, and by considering only gene codons with a minimum coverage cut-off of 20X across all metagenomes of interest. Controlling the minimum coverage of gene codons across metagenomes improves confidence in variability analyses. Table S5 reports the SAAV density values for all core S-LLPA genes.

### SAAV coordinates and the frequency of amino acid substitution (AAS) types

Each SAAV was defined by a gene codon coordinate, an AAST (two most frequent amino acids), and a metagenome. Anvi’o attributed BLOSUM62 and BLOSUM90 values (ranging from ‘-6’ to ‘+3’) to each AAST.

### Application of Deep Learning to SAAV data

To estimate an unbiased distance between our metagenomes based on SAAVs, we used a novel deep neural network modification called the auto-clustering output layer (ACOL). Briefly, ACOL relies on a recently introduced graph-based activity regularization (GAR) technique for competitive learning from hyper-dimensional data to demarcate fine clusters within user-defined ‘parent’ classes [44]. In this application of ACOL, however, we modified the algorithm so it can reveal latent groups in our SAAVs in a fully unsupervised manner through frequent random sampling of SAAVs to create pseudo-parent class labels instead of user-defined classes [45]. See the Supplementary Methods for the details of pseudo parent-class generation, and distance estimation.

### Computing the SAAV to SNV ratio

For a genome with *N* genes, let 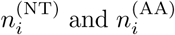 be the number of SNVs and SAAVs in gene *i*, respectively. We defined *S*, the SAAV to SNV ratio, as the average ratio across all genes of 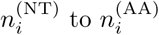:

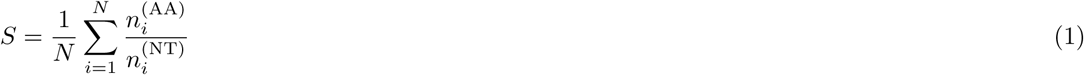

Since each SAAV corresponds to at least one SNV, ranges from 0 to 1, where 0 and 1 correspond to completely synonymous and non-synonymous variation, respectively. Although the ratio of non-synonymous to synonymous nucleotide substitutions (denoted *K_a_*/*K_s_* or *d_N_*/*d_S_*) is commonly used to quantify selective pressure in bacterial populations [46, 47], this single-nucleotide approach is invalid for the S-LLPA population, since on average 22.5% of SNVs per metagenome co-occurred with other SNVs in the same codon. Hence, we used as an alternative proxy for selective pressure that bears qualitative similarity to and is generalizable to populations containing dense variation, such as S-LLPA.

### 3D structure of proteins using template-based protein structure modeling

We used a template-based structure modeling tool, RaptorX Structure Prediction [48], to predict structures of S-LLPA amino acid sequences based on available data from the Protein Data Bank (PDB) citebernstein1977. We used blastp in NCBI’s BLAST distribution to identify core S-LLPA genes that matched to an entry with at least 30% similarity over the length of the given core gene. We then programmatically mapped SAAVs from metagenomes onto the predicted tertiary structures, and used PyMOL [49] to visualize these data. We colored SAAVs based on RaptorX-predicted structural properties, including solvent accessibility and secondary structure.

We used the aov function in R to perform one way ANOVA tests, used the ggplot2 [50] package for R to visualize the relative distribution of S-LLPA genes and geographic distribution of proteotypes, and finalized all figures using an open-source vector graphics editor, Inkscape (available from http://inkscape.org/).

## Data availability

The TARA Oceans and Ocean Sampling Day metagenomes are publicly available through the European Bioinformatics Institute (accession IDs ERP001736, and ERP001736, respectively). We also made available (1) SAR11 isolate genomes (doi:10.6084/m9.figshare.5248945.v1), (2) the anvi’o contigs database and merged profile for SAR11 genomes across metagenomes (doi:10.5281/zenodo.835218) and the static HTML summary for the mapping results (doi:10.6084/m9.figshare.5248453), (3) the SAR11 metapangenome (doi:10.6084/m9.figshare.5248459), single-nucleotide and single-amino acid variant reports for S-LLPA across 74 TARA Oceans metagenomes (doi:10.6084/m9.figshare.5248447), and (4) SAAVs overlaid on predicted tertiary structures of 58 core S-LLPA genes (doi:10.6084/m9.figshare.5248432). The URL https://anvi-server.org/merenlab/sar1Dmetapangenome serves an interactive version of the SAR11 metapangenome, and the URL http://anvio.org/data/S-LLPA-SAAVs/ serves an interactive web page to investigate the link between SAAVs and predicted protein structures.

## Supplementary Information

Characterization of the core S-LLPA genes: We first identified 78 metagenomes in which the mean coverage of HIMB83 was >50X. Despite the systematic low occurrence of certain genes, a large fraction of the HIMB83 gene pool (n=1,470) was detected within this set of 78 metagenomes (Figure S2). In order to determine the core S-LLPA genes, we first disregarded four metagenomes (including three from the Mediterranean Sea) that displayed an unusually high coefficient of gene coverage variation (Figure 2, panel A), a factor indicative of non-specific read recruitments originating from other abundant populations. Using the final selection of 74 metagenomes, we then removed genes exhibiting a mean coverage value five times below (n=630) or above (n=46) the average mean coverage of all genes in at least one metagenome. Our analysis resulted in the identification of 799 core genes containing 757,000 nucleotides that we considered maintained a sufficient linkage with S-LLPA regardless of geographic distances.

## Supplementary Methods

Automatic Learning of Latent Annotations on Neural Networks with Auto-clustering Output Layer using Pseudo Parent-Classes

Our approach builds upon the previous study on learning of latent annotations on neural networks using auto-clustering output layer (ACOL) when a coarse level of supervision is available for all observations, i.e. parent-class labels, but the model has to learn a deeper level of latent annotations, i.e. sub-classes, under each one of parents [45]. ACOL is a novel output layer modification for deep neural networks to allow simultaneous supervised classification (per provided parent-classes) and unsupervised clustering (within each parent) where clustering is performed with a Graph-based Activity Regularization (GAR) technique recently proposed in [44]. More specifically, as ACOL duplicates the softmax nodes at the output layer for each class, GAR allows for competitive learning between these duplicates on a traditional error-correction learning framework.

To investigate clusters of metagenomes with respect to the SAAVs found in core S-LLPA genes, we modified ACOL to learn latent annotations in a fully unsupervised setup by substituting the real, yet unavailable, parent-class information with a pseudo one, i.e. randomly generated pseudo parent-classes. To generate examples for a pseudo parent-class, we choose a domain specific transformation to be applied to every sample in the dataset. The transformed dataset constitutes the examples of that pseudo parent-class and every new transformation - for this particular case the random sampling of codon positions - generates a new pseudo parent-class. Naturally, the main classification task performed over these pseudo parent-classes does not represent any meaningful knowledge about the data by itself. However, frequent and random selection of these pseudo parent-classes allow the ACOL neural network to learn sub-classes of these pseudo parents without bias. While each sub-class corresponds to a latent annotation which may or may not be meaningful, the combination of these annotations learned through abundant and concurrent clusterings reveals an unbiased and robust similarity metric between different metagenomes.

Pseudo parent-class generation and similarity metric calculation are described thoroughly in the following subsections (or appendices) along with the details of model training in which we apply a different unsupervised regularization rather than adopting GAR.

### Pseudo parent-class generation

Consider an *n_s_* × *n_p_* × *n_f_* metagenomics dataset represented by the 3-D tensor **D**, where *n_s_* is the number of metagenomes to be clustered, *n_p_* is the number of codon positions per metagenome and *n_f_* is the number of features representing each codon position. Specifically, our dataset can be specified by a 74 × 37,416 × 2 tensor, as each codon position is represented by the two most frequent amino acids found in this position.

To generate the examples of *i*^th^ pseudo parent-class, we randomly sample *n_ṕ_* positions out of *n_p_* with replacement. Resulting *n_s_* × *n_ṕ_* × *n_f_* subsets which correspond to *d* = *n_ṕ_n_f_* dimensional *n_s_* examples of pseudo parent-class *i*. This procedure is repeated for *n_ψ_* times, and ultimately we obtain an *m* × *d* input matrix ***X*** = [***x***_1_,…, ***x***_*m*_]^*T*^ and corresponding pseudo parent-class labels ***t*** = [*t*_1_,…, *t_m_*]^*T*^ to train a neural network, where *n_ψ_* is the number of the pseudo parents and m is the total number of the examples generated in this procedure, such that *m* = *n_ψ_n_s_*.

We select *n_ṕ_* = 2000 and *n_ψ_* = 1000 to produce the results in this article, therefore ***X*** is 74000 × 4000 matrix where each one of 1000 pseudo parents is equally represented by 74 examples sampled from different metagenomes. We also keep track of the metagenome labels ***q*** = [*q*_1_,…, *q_m_*]^*T*^, indicating the source metagenome of every example. This information will be used later to produce the similarity matrix.

Algorithm 1 below describes the entire sampling and pseudo parent-class generation procedure.

#### Algorithm 1

Pseudo-class generation

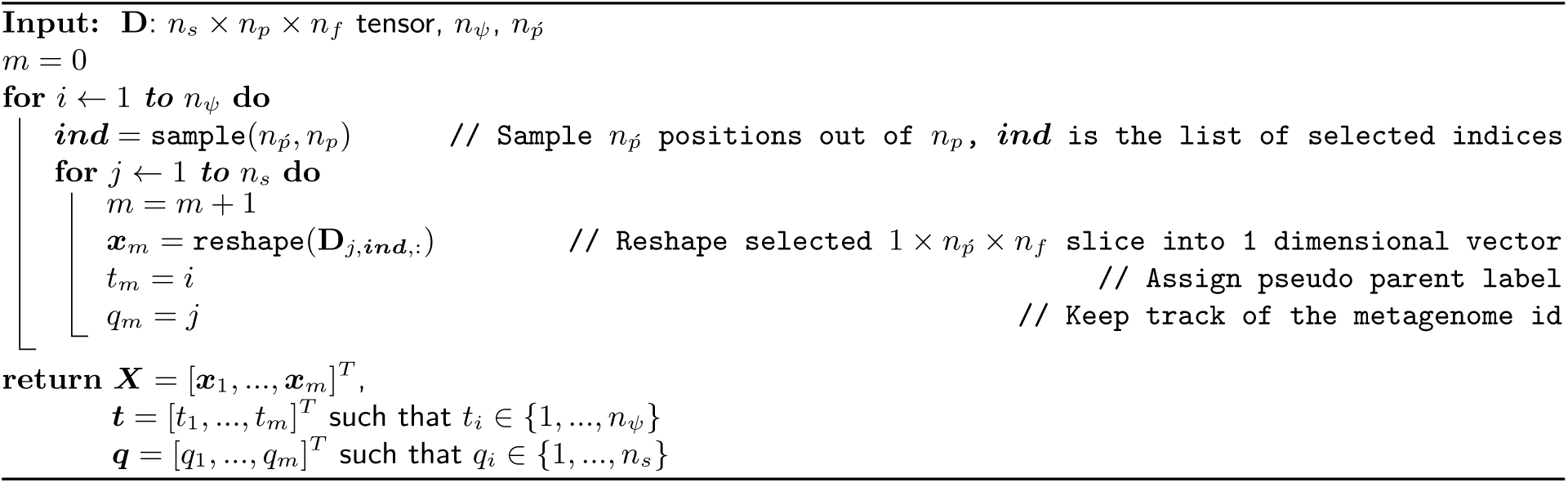

### Learning Latent Annotations and Constructing the Similarity Matrix

Neural networks define a family of functions parameterized by weights and biases which define the relation between inputs and outputs. In multi-class categorization tasks, outputs correspond to class labels, hence in a typical output layer structure there exists an individual output node for each class. An activation function, such as softmax is then used to calculate normalized exponentials to convert the previous hidden layer’s activities, i.e. scores, into probabilities.

Unlike traditional output layer structure, ACOL defines more than one softmax node (*k* duplicates) per class (in this particular case, we prefer to use *pseudo parent*-*class* term to emphasize that these classes are not expert-defined but automatically generated depending on random sampling). Outputs of *k* duplicated softmax nodes that belong to the same pseudo parent are then combined in a subsequent pooling layer for the final prediction. Training is performed in the configuration shown in Figure 4. This might look like a classifier with redundant softmax nodes. However, duplicated softmax nodes of each pseudo parent are specialized due to dropout[51] and the unsupervised regularization applied throughout the training in a way that each one of *n* = *n_ψ_k* softmax nodes represents an individual sub-class of a pseudo parent, i.e. latent annotation.

**Figure 4.**
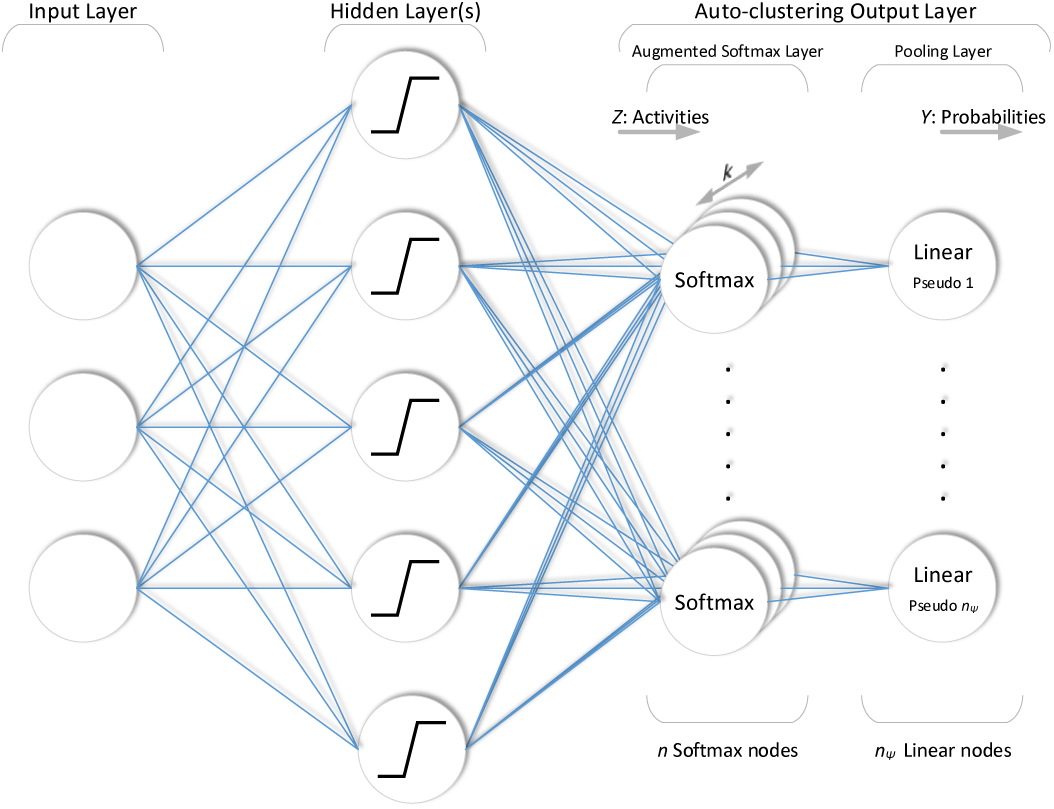
Neural network structure with the ACOL. Each softmax node corresponds to an individual sub-class of a pseudo parent, i.e. latent annotation. During feedforward operation of the network, pooling layer calculates final pseudo parent-class predictions through sub-class probabilities.

Consider a neural network with ACOL consisting of *L* – 1 hidden layers where *l* denotes the individual index for each layer such that *l* ∈ {0,…, *L*}. Let ***Y***^(*l*)^ denote the output of the nodes at layer *l*. ***Y***^(0)^ = ***X*** is the input and *f*(***X***) = *f*^(*L*)^(***X***) = ***Y***^(*L*)^ = ***Y*** is the output of the entire network. ***W***^(*1*)^ and ***b***^(*1*)^ are the weights and biases of layer *l*, respectively. Then, the feedforward operation of the neural network can be written as

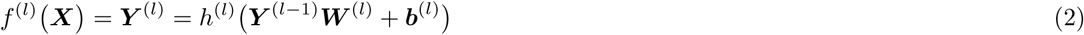

where *h*^(*1*)^(.) is the activation function applied at layer *l*. For the sake of clarity, let us specify the activities going into the augmented softmax layer of this network such that

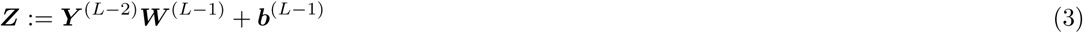

and its positive part as ***B*** such that

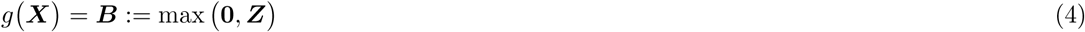

both of which correspond to *m* × *n* matrices, where *n* = *n_ψ_k* and *k* is the clustering coefficient of ACOL and chosen as 4 for this application. Then, the output of a neural network with ACOL can be written in terms of ***Z*** as

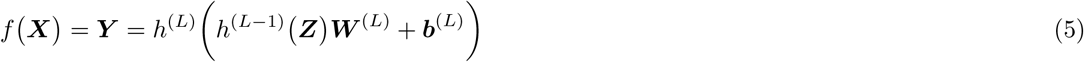

where ***Y*** is an *m* × *n_ψ_* matrix in which *Y_ij_* is the probability of the *i*^th^ example belonging to the *j*^th^ pseudo parent. Since *h*^(*L*–1)^(.) and *h*^(*L*)^(.) respectively correspond to softmax and linear activation functions and ***b***^(*L*)^ = **0**, then (5) further simplifies into

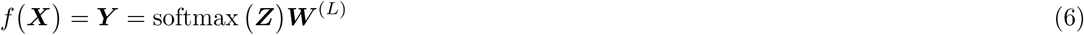

where ***W***^(*L*)^ is an *n* × *n_ψ_* matrix representing the constant weights between augmented softmax layer and pooling layer.

Also, unsupervised regularization 𝓤(.) is applied to ***B*** to penalize the range of its distribution. The overall objective function of the training then becomes

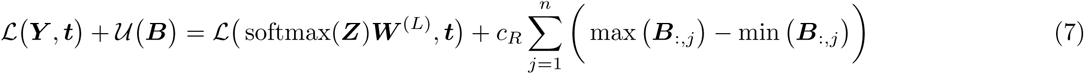

where 𝓛(.) is the supervised log loss function, ***t*** = [*t*_1_,…, *t_m_*]^*T*^ is the vector of the provided pseudo parent labels such that *t*_1_ ∈ {1,…, *n_ψ_*}, ***B***_:,*j*_ corresponds to *j*^th^ column vector of matrix ***B*** and *c_R_* is the weighting coefficient.

Training of the proposed framework is performed according to simultaneous supervised and unsupervised updates resulting from the objective function given in (7). We adopt stochastic gradient descent in the mini-batch mode [52] for optimization. Algorithm 2 below describes the entire training procedure. Along with unsupervised regularization, dropout method [51] is also applied to the augmented softmax layer to distribute the examples across the duplicated softmax nodes. For each mini-batch, an 1 × *n* row vector ***r*** of independent Bernoulli random variables is sampled and multiplied element-wise with the output of the augmented softmax layer. This operation corresponds to dropping out each one of the *n* softmax nodes with the probability of *p* for that batch.

#### Algorithm 2

Model training

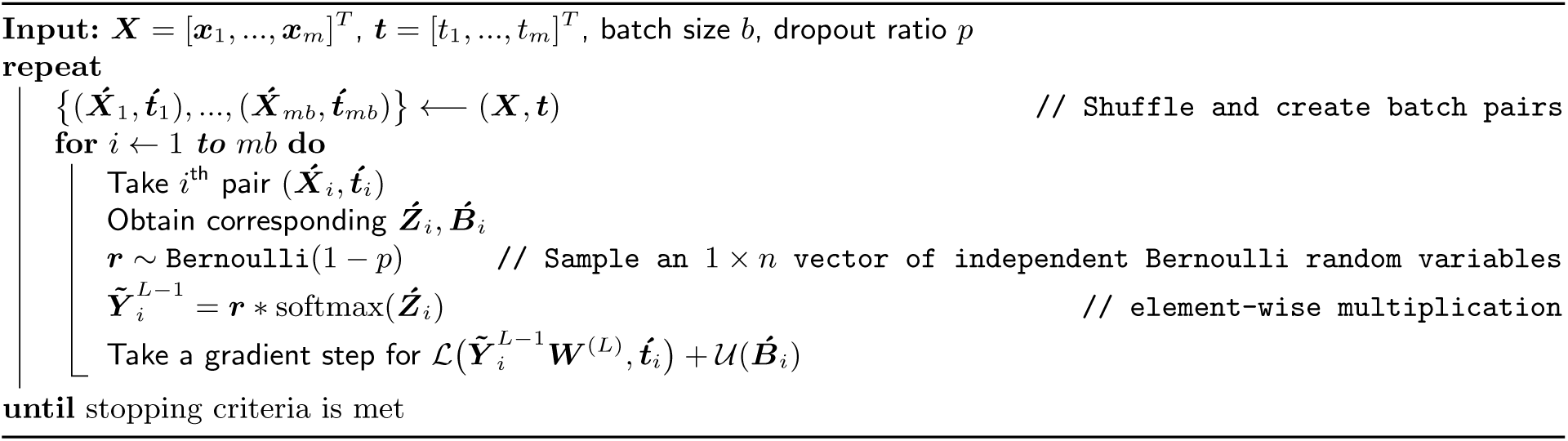

After training phase is completed, network is simply truncated by completely disconnecting the pooling layer as shown in Figure 5 and the rest of the network with trained weights is used to assign the annotations to each example. This assignment can be described as

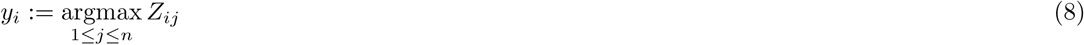

where *y_i_* is the annotation assigned to *i*^th^ example. Using the assigned annotations ***y*** and the metagenome labels ***q***, the similarity matrix ***S*** used to obtain the dendrogram provided in this article is constructed as shown in Algorithm 3. Since applying dropout to distribute the examples across the duplicated softmax nodes also introduces a variance to the assigned annotations, we take the average of the similarity matrices obtained during the last 20 epochs of the training. Moreover, to reduce the variance due to the random generation of the pseudo parent-classes, we repeat the entire procedure 100 times with a new selection of pseudo parents for each repetition.

**Figure 5.**
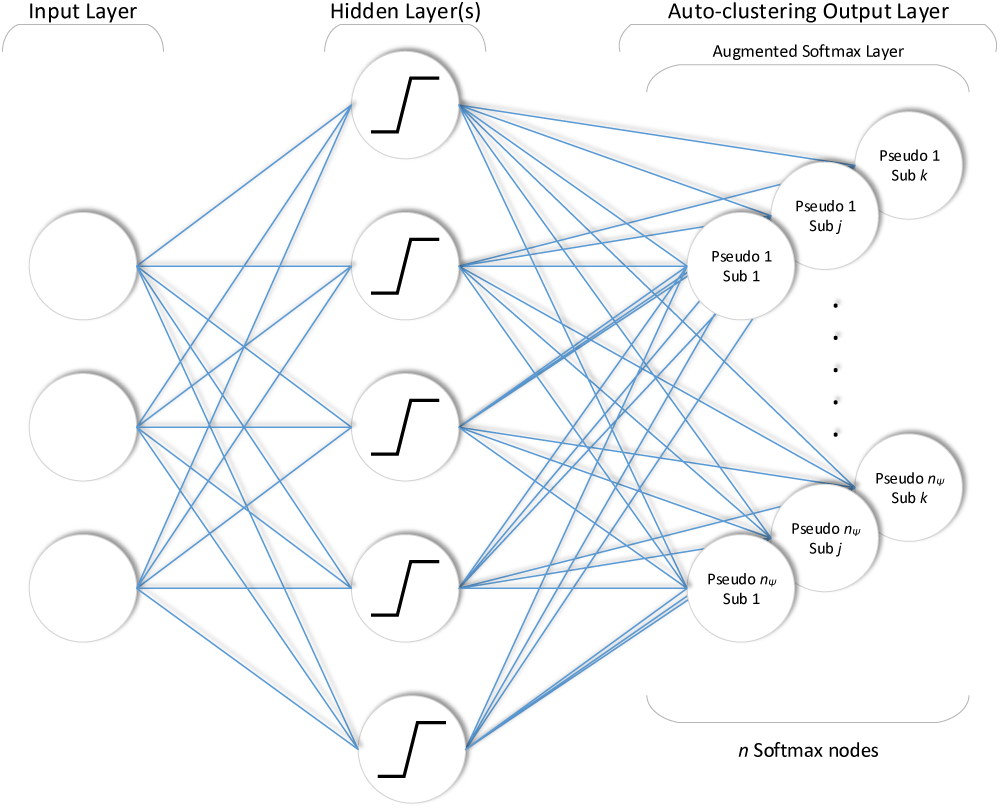
After training the pooling layer is simply disconnected from rest of the network. The rest of the network with trained weights is used to obtain the assigned annotations. This operation can be described as *y_i_*: = argmax_1<*j*<*n*_ *Z_ij_*.

#### Algorithm 3

Obtaining similarity matrix

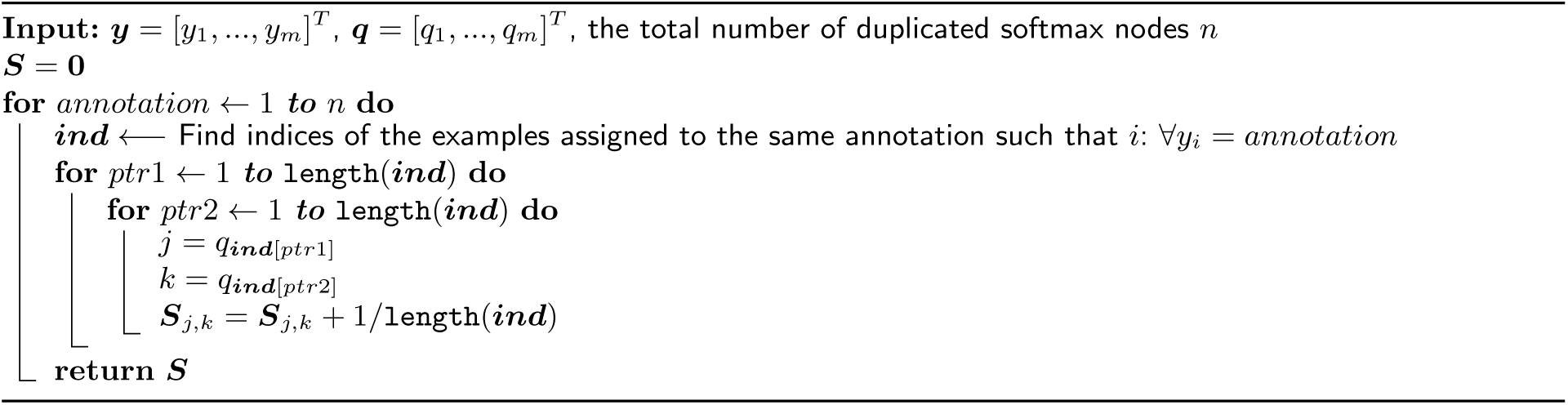

## Author contributions

T.O.D and A.M.E conceived the study, and wrote the paper. T.O.D, E.K, O.C.E, and A.M.E developed analysis tools. O.K and I.U developed the ACOL algorithm. All authors contributed to the analysis of the data, and reviewed and revised the drafts of the paper.

## Acknowledgements

We thank the TARA Oceans consortium for generating metagenomic datasets of great legacy, as well as all researchers involved in the characterization of SAR11 isolates. We thank Edward Delong, Mike Lee, and Alon Shaiber for helpful discussions. This study was supported by the Frank R. Lillie Research Innovation Award and the start-up funds from the University of Chicago to AME.

## Competing interests

The authors declare that they have no competing interests.

